# Detection and removal of barcode swapping in single-cell RNA-seq data

**DOI:** 10.1101/177048

**Authors:** Jonathan A. Griffiths, Arianne C. Richard, Karsten Bach, Aaron T.L. Lun, John C Marioni

## Abstract

Barcode swapping results in the mislabeling of sequencing reads between multiplexed samples on the new patterned flow cell Illumina sequencing machines. This may compromise the validity of numerous genomic assays, especially for single-cell studies where many samples are routinely multiplexed together. The severity and consequences of barcode swapping for single-cell transcriptomic studies remain poorly understood. We have used two statistical approaches to robustly quantify the fraction of swapped reads in each of two plate-based single-cell RNA sequencing datasets. We found that approximately 2.5% of reads were mislabeled between samples on the HiSeq 4000 machine, which is lower than previous reports. We observed no correlation between the swapped fraction of reads and the concentration of free barcode across plates. Furthermore, we have demonstrated that barcode swapping may generate complex but artefactual cell libraries in droplet-based single-cell RNA sequencing studies. To eliminate these artefacts, we have developed an algorithm to exclude individual molecules that have swapped between samples in 10X Genomics experiments, exploiting the combinatorial complexity present in the data. This permits the continued use of cutting-edge sequencing machines for droplet-based experiments while avoiding the confounding effects of barcode swapping.

## Introduction

Recent reports have shown that the DNA barcodes used to label multiplexed libraries can “swap” on patterned flow-cell Illumina sequencing machines, including the HiSeq 4000, HiSeq X, and NovaSeq (Sinha *et al.*, 2017; Costello *et al.*, 2017). This results in mislabeling whereby reads assigned to one sample derive from molecules in another, thus compromising the interpretation of many-omic assays (Figure 1). Barcode swapping is particularly problematic for single-cell RNA sequencing (scRNA-seq) experiments, where many libraries are routinely multiplexed together. For example, barcode swapping could lead to cells that appear to falsely express particular marker genes, or yield spurious correlation patterns that may confound clustering and other analyses.

**Figure 1:**
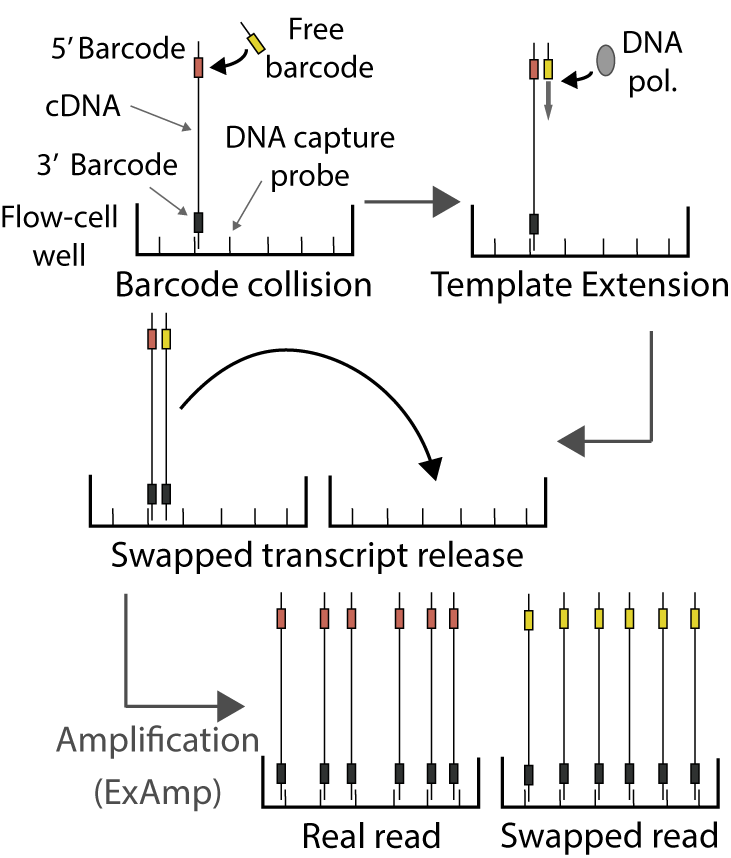
A schematic of the mechanism for barcode swapping, as proposed by Sinha *et al.* (2017). On new models of the Illumina sequencing machines, flow cell seeding and DNA amplification take place simultaneously, without any washes of the flow cell between steps. As a result, free sample indexing barcodes remain in solution and can be inadvertently extended using DNA molecules from libraries with different barcodes as templates. The transfer of mislabeled molecules between nanowells of the flow cell results in clustering and sequencing of artefactually labelled DNA molecules.

The severity and consequences of barcode swapping in scRNA-seq studies remain poorly understood. Sinha *et al.* (2017) estimated swapping rates of “up to 5-10%” from a plate-based scRNA-seq experiment; however, these estimates were obtained from only two wells in a single micro-well plate. The lack of replication makes it difficult to generalize the results to other scRNA-seq studies. Furthermore, the effect of barcode swapping on high-throughput droplet-based scRNA-seq protocols (Zheng *et al.*, 2017) has not been explored. This is a key consideration due to the increasing use of droplet-based methods for large-scale single-cell studies (Schiebinger *et al.*, 2017; Dixit *et al.*, 2016) where many samples are necessarily multiplexed together for efficient sequencing.

Here, we robustly quantify the fraction of swapped reads in each of two plate-based single-cell RNA sequencing datasets. We found that approximately 2.5% of reads were mislabeled between samples on HiSeq 4000, and observed no correlation between the swapped fraction of reads and the concentration of free barcode across plates. Furthermore, we demonstrate that barcode swapping can generate complex but artefactual cell libraries in droplet-based scRNA-seq data. To eliminate these artefacts, we developed a computational method to exclude swapped reads in 10X Genomics experiments, enabling the continued use of cutting-edge sequencing machines for droplet-based experiments.

## Barcode swapping in plate-based single cell RNA-seq experiments

A number of widely-used scRNA-seq library preparation methods isolate and process individual cells in wells of a microwell plate before performing library preparation in parallel (Picelli *et al.*, 2014; Jaitin *et al.*, 2014; Hashimshony *et al.*, 2016). A unique combination of sample barcodes characterises the library associated with each cell, usually by adding a different barcode to each end of a cDNA molecule. One barcode typically indexes the row position for each cell on the microwell plate, while the other barcode indexes the column position. Swapping of either or both barcodes therefore moves reads between cell libraries. We used two independent plate-based scRNA-seq datasets to quantify the swapping fraction, i.e., the fraction of all cDNA reads across all sequencing libraries multiplexed on a single flow cell lane that were mislabelled.

In the first dataset (referred to as the “Richard dataset”, see Supplementary File Section 3), two plates of single mouse T cells were multiplexed for sequencing on a HiSeq 4000 instrument. Each of these plates used entirely different sets of column and row barcodes: none of the barcodes were reused between the plates (Figure 2A). As such, there are a set of barcode combinations that should contain zero reads (“impossible” combinations), as the two sets of barcodes for these combinations were never mixed during the experiment. However, reads mappable to the mouse genome were still present in the impossible combinations at approximately 1% of the frequency in the expected combinations (Figure 2B). This cannot be explained by contamination from free-floating nucleic acids, which can only affect the expected combinations used during library preparation. Indeed, the number of reads in each impossible combination was proportional to the number of reads in the real cell libraries that shared exactly one barcode with the impossible combination (Figure 2C). This is consistent with read misassignment to an impossible combination, due to swapping of a single barcode from the pool of cDNA in the expected combinations.

**Figure 2:**
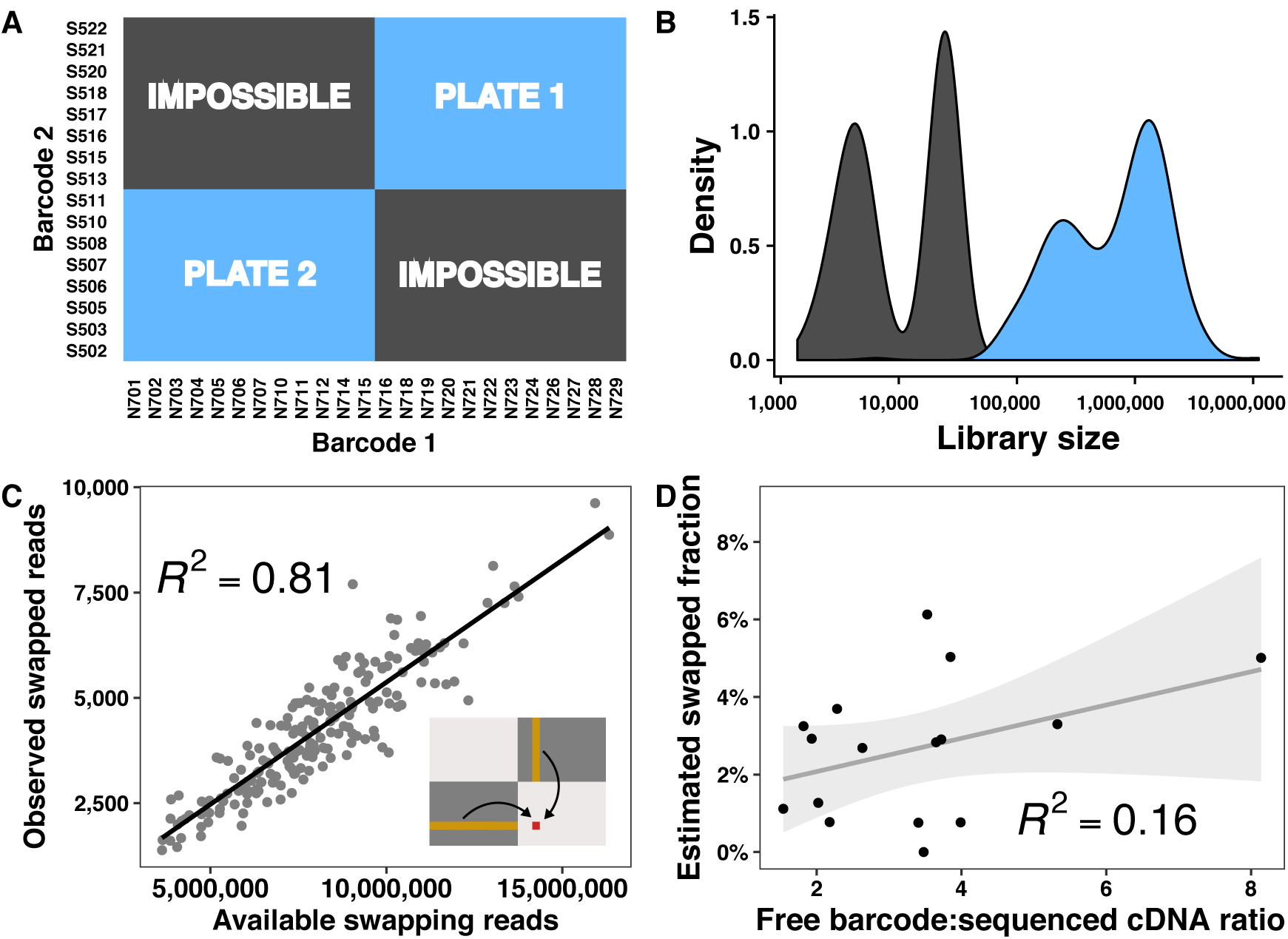
Characterization of barcode swapping in plate-based scRNA-seq experiments. (**A**) The experimental design of the Richard dataset. Two 96-well plates of cells were multiplexed for sequencing. Expected barcode combinations are marked in blue, while impossible barcode combinations are marked in grey. (**B**) Distribution of the library sizes (i.e., number of mapped reads) in the expected and impossible barcode combinations. (**C**) Library size of each impossible combination (observed swapped reads), plotted against the sum of the library sizes of the expected combinations that share exactly one barcode with that impossible combination (available swapped reads). An example is illustrated graphically in the inset Figure for one impossible combination (red) and the contributing expected combinations (orange). The gradient represents the fraction of available reads from the expected combinations that swap into each impossible combination. (**D**) Estimated swapping fractions for different plates of the Nestorowa *et al.* (2016) dataset, plotted against the ratio of the concentration of free barcode to the concentration of cDNA of the correct length for sequencing. A linear regression fit is shown along with its 95% confidence interval. The slope of the fitted line is not significantly different from 0 (*p* = 0.129).

To estimate the swapped fraction in the Richard dataset, we regressed the library size of the impossible combinations against the summed library sizes of the real cells that shared exactly one barcode (see Supplementary File Section 3 for more details). This yielded an estimate of the swapped read fraction of 2.18*±*0.08%. For comparison, we repeated this procedure on the same libraries sequenced on HiSeq 2500, yielding a much lower swapped fraction estimate of 0.22*±*0.01%. This is consistent with the proposed mechanism of barcode swapping on the new Illumina machines. Notably, our estimates are calculated over many wells with impossible combinations, offer an estimate of uncertainty, and are robust to contamination. This represents an improvement over previous estimates (Sinha *et al.*, 2017), which only sought to technically demonstrate the existence of swapping by considering two wells of a single plate.

In the second dataset (Nestorowa *et al.* (2016), see Supplementary File Section 4), we considered plates of single cells whose libraries had been sequenced on both the HiSeq 2500 and 4000. We modelled each cell’s gene expression levels in the HiSeq 4000 data as a linear combination of contributions from the HiSeq 2500 data. Specifically, for each cell, we considered contributions from itself, cells that share exactly one barcode, and cells that share no barcodes (see Supplementary File Section 4 for details). The relative contribution from other cells was used to estimate the swapping fraction across all cells in the plate. Across 16 independent plates of single cells (each of which was sequenced separately), we estimated a range of swapped read fractions with mean 2.275*±*0.359%, consistent with the first experiment (Figure 2D). We also observed that nearly all expressed genes were affected by swapping (Supplementary File Section 4.5), consistent with its global effects on the pool of sequencable DNA.

Given the range of estimated swapping fractions in the Nestorowa *et al.* (2016) data, we reasoned that we could identify the factors driving barcode swapping by considering the library characteristics of each plate. Specifically, we investigated the association with the amount of free library barcode, which has previously been linked to swapping rates (Sinha *et al.*, 2017). We used Agilent’s Bioanalyzer Expert 2100 software to quantify the amount of free barcode (DNA lengths 45-70bp) and the amount of sequencable cDNA (400-800bp) in the multiplexed library from each plate. However, we did not observe a strong correlation of swapping fraction estimates with the ratio of free barcode to sequencable cDNA (Figure 2D). Similarly, no correlation was observed with the total amount of free barcode per plate, or the ratio of free barcode concentration to mappable reads (Supplementary File Section 4.4). This suggests that the extent of barcode swapping is not primarily determined by the amount of free barcode, at least not in experiments using typical barcode concentrations.

## Removing the effects of barcode swapping in plate-based experiments

The most obvious solution for barcode swapping is to use a sequencing machine that does not use a patterned flow cell, which we have shown reduces the swapping rate by an order of magnitude. Where this is not possible, an approach for computationally “unmixing” the expression profiles has been described (Larsson *et al.*, 2017), although the user must specify the swapping fraction for accurate correction. Unfortunately, our results indicate that the swapping rate varies across plates of a scRNA-seq experiment (Figure 2D), so it is unlikely that a single estimate of the swapping fraction is appropriate for all experiments.

For general plate-based scRNA-seq experiments, we recommend the use of an experimental design similar to that in the Richard dataset (Figure 2A). By leaving a fraction of possible barcode combinations unoccupied, a researcher can robustly estimate the swapped fraction of reads. This serves as a useful quality control metric for individual experiments, whereby datasets with high swapping rates can be flagged and discarded to avoid generating misleading biological results. The swapping fraction estimate for each experiment can also be used to improve the accuracy of any computational correction (Larsson *et al.*, 2017).

Unique dual indexing represents another experimental solution to barcode swapping (Costello *et al.*, 2017). Under this scheme, two unique barcodes are used for each sample in a multiplexed sequencing experiment. A single barcode swap will move reads to barcode combinations that are not used by any other sample, thus avoiding any mixing of libraries between samples. However, the need for unique indexes greatly restricts the number of libraries that can be multiplexed for a given number of barcodes (see Supplementary File Section 7 for scalability calculations). This is particularly problematic for single-cell studies where large numbers of cell libraries need to be multiplexed for efficient sequencing. To use unique dual indexing in such cases, a researcher must have a large number of available barcodes, which may not be practical.

## Barcode swapping in droplet-based single-cell RNA-seq experiments

New single-cell RNA-seq protocols use microfluidic systems to massively multiplex library preparation by capturing individual cells in droplets (Macosko *et al.*, 2015; Zheng *et al.*, 2017). These methods enable the efficient generation of thousands of single-cell libraries in a single experiment. Cell labelling is achieved by the incorporation of a cell barcode in the reverse transcription step that occurs in each droplet. Each cell barcode is selected randomly from a large pool of possible sequences. A single sample barcode is then used to label different batches of single cells for multiplexed sequencing. The cell barcode is never free in solution; only the sample barcode is expected to swap.

We consider two major effects of barcode swapping in droplet-based experiments. Firstly, it is possible that the same cell barcode is used in two or more multiplexed samples. Between these samples, swapping will mix transcriptomes of different cells labelled with the same cell barcode, similar to the effect observed in plate-based assays. The second effect arises when a “donor” sample contains a cell barcode that is not present in another “recipient” sample. Swapping of molecules labelled with this donor-only barcode will produce a new artefactual cell library in the recipient sample. This new library will have a similar expression profile to the original cell in the donor sample and may be identified as a real cell. Indeed, swapping from cell libraries that are especially large may generate artefactual libraries in recipient samples that are as large and complex as real cells, making it difficult to find and remove them.

We demonstrated the existence of artefactual cell libraries in real data by testing whether samples from droplet-based experiments shared more cell barcodes than expected by chance. We obtained 10X Genomics data for human breast tumour cells and mouse epithelial cells, sequenced separately on the HiSeq 4000. In both of these experiments, at least one sample comparison exhibited excess sharing according to a hypergeometric test (Supplementary Figures 29-30, Supplementary File Section 5.2). We also obtained 10X Genomics data for mouse embryonic cells sequenced on a HiSeq 2500, and resequenced the aforementioned mouse epithelial cells on the HiSeq 2500. In both of these experiments, no excess sharing was observed (Supplementary Figures 31-32). This is again consistent with barcode swapping on the new Illumina machines.

## Removing the effects of barcode swapping

One obvious solution to mitigate the effect of barcode swapping is to discard any cell libraries with shared cell barcodes across multiplexed samples. This removes both homogenised and swap-derived artefactual libraries from further analysis, thus avoiding misleading conslusions driven by barcode swapping. However, cell-based removal is not appropriate when many cells are captured across many samples for a single multiplexed sequencing run. This is because many cells will share cell barcodes by chance, even in the absence of barcode swapping. Removal of these cells will result in unnecessary loss of data (Figure 3A). For example, applying this strategy to 30 multiplexed samples of 20,000 cells each would exclude over 50% of cell libraries. An alternative approach is necessary for high-throughput droplet scRNA-seq datasets that are now routinely generated (Stoeckius *et al.*, 2017; Schiebinger *et al.*, 2017; Dixit *et al.*, 2016).

**Figure 3:**
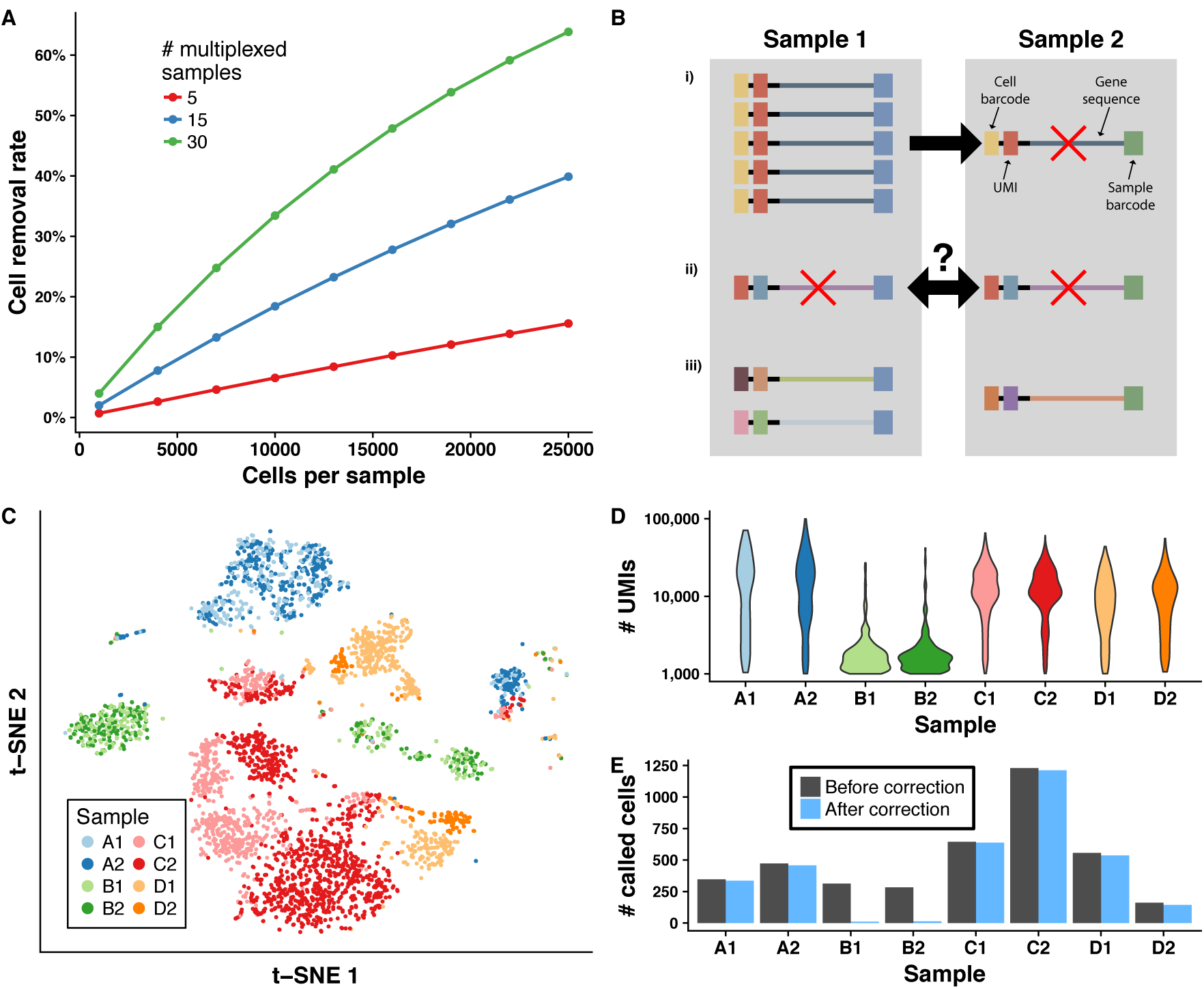
Characterization of barcode swapping in droplet-based scRNA-seq experiments. (**A**) The expected number of cells with shared cell barcodes in 10X samples that have been multiplexed for sequencing, for different numbers of samples and different numbers of captured cells per sample. (**B**) A schematic of our method to remove swapped reads from droplet data. Reads found in different samples with the same combination of UMI, cell barcode, and aligned gene were considered to have swapped. If most reads (≥ 80%) were present in one sample, we excluded the molecule from all other samples (i). If reads were more evenly spread across samples, we excluded the molecule from all samples (ii). Reads in one sample only were retained (iii). (**C**) *t*-SNE plot of the expression profiles of mouse epithelial cells (Maaten & Hinton, 2008). Each point represents a cell that is coloured by sample. Letters correspond to different experimental conditions while numbers represent biological replicates. (**D**) The distribution of the library sizes for called cells in each sample. Cells were called using *emptyDrops* (Lun, 2018), with an FDR threshold of 1% and a minimum of 1,000 UMIs. (**E**) The number of called cells for each sample, before and after application of our swapped read exclusion algorithm.

We have developed a computational method that removes individual swapped reads from 10X Genomics data, avoiding the exclusion of entire cell libraries. Specifically, we considered molecules across multiplexed samples that contain the same combination of unique molecular identifier, cell barcode, and aligned gene. Due to combinatorial complexity, these molecules are extremely unlikely to arise by chance, and are almost certain to be generated by barcode swapping. For each observed combination of these labels, we calculated the fraction of reads that were present in each sample. Where one sample contained the majority of all reads for a molecule (≥ 80%), we considered this as the sample-of-origin, and removed the molecule count from all other samples. Where this was not the case, we removed the molecule from all samples (Figure 3B), as an unambiguous determination of the sample-of-origin was not possible.

We applied our method to the aforementioned mouse epithelial cell dataset sequenced on the HiSeq 4000. In one experiment, two samples (B1, B2) appeared to contain many cells with expression profiles that were distinct from those of the cells in the other samples (Figure 3C). We observed that cells in these samples had smaller library sizes than in other samples (Figure 3D). Further inspection revealed that many cell barcodes in B1 and B2 were also present in other samples (Supplementary File Section 5.3). We hypothesised that the majority of cell libraries in samples B1 and B2 derived from barcode swapping. Exclusion of swapped reads resulted in the loss of nearly all called cells from these samples (Figure 3E, Supplementary File Section 6.2.3), indicating that they consisted almost entirely of artefactual swapped libraries.

These results demonstrate the importance of excluding swapped reads prior to further analysis. Failure to do so would have resulted in misleading biological conclusions if the artefactual cells were used in analyses such as clustering and detection of differentially expressed genes. Indeed, the artefactual cells form their own cluster in Figure 3C, and could be misinterpreted as a cell type exclusive to samples B1 and B2. We also observed that 2.5% of cell libraries from the other samples were no longer called as cells after removal of swapped reads. While this represents the removal of a smaller number of artefactual cells, it may be important in studies involving rare cell types where the presence of a few cells can affect the interpretation of the results.

As a control, we applied our method to two 10X Genomics experiments using mouse epithelial cells that were not multiplexed together. Here, our method removed a negligible number of molecules (*<*0.0005%, Supplementary File Section 6.2.4). This demonstrates that our method is able to specifically exclude swapped reads. Our method is implemented in the *DropletUtils* Bioconductor package for 10X Genomics experiments, and can be easily applied in a conventional analysis pipeline.

## Conclusion

Using plate-based scRNA-seq datasets, we have reproducibly estimated the fraction of barcode-swapped reads on the HiSeq 4000 to be approximately 2.5%, which is lower than previously reported (Sinha *et al.*, 2017). Different amounts of free DNA barcode did not affect our swapping fraction estimates, suggesting that free barcode concentration is not the primary factor determining the variation in barcode swapping rates across experiments. We recommend that plate-based scRNA-seq experiments that reuse cell barcodes should continue to be sequenced on non-patterned flow-cell machines such as the HiSeq 2500 to minimise barcode swapping. We have also implemented a compuational method for removing swapped reads from 10X Genomics data without removing entire cell libraries. This permits the cost-effective use of the highest-throughput sequencing machines (e.g., the HiSeq 4000) for large-scale droplet scRNA-seq experiments while avoiding the confounding effects of barcode swapping.

## DATA AVAILABILITY STATEMENT

A Github repository (https://github.com/MarioniLab/BarcodeSwapping2017) contains a detailed report that expands on the analyses described herein, describing models and showing results. The repository also contains a script to download the data, and the code used to generate the report.

## ACKNOWLEDGMENTS

We thank Gillian Griffiths, Berthold Göttgens, Fernando Calero, Sonia Nestorowa, Fiona Hamey, Carlos Caldas, Dimitra Georgopoulou, and Alejandro Bruna for help with data access.

## AUTHOR CONTRIBUTIONS

J.A.G. performed data analyses; A.T.L.L. and K.B. contributed code; K.B. and A.C.R. generated data; J.A.G., A.T.L.L., and J.C.M. wrote the manuscript; all authors read and approved the final manuscript.

